# A high-content screen identified ingenol-3-angelate as an enhancer of B7-H3-CAR T cell activity by increasing B7-H3 protein expression on the target cell surface via PKCα activation

**DOI:** 10.1101/2023.05.26.542130

**Authors:** Ha Won Lee, Carla O’Reilly, Alex N. Beckett, Duane G. Currier, Taosheng Chen, Christopher DeRenzo

## Abstract

CAR T cell therapy is a promising approach to improve outcomes and decrease toxicities for patients with cancer. While extraordinary success has been achieved using CAR T cells to treat patients with CD19-positive malignancies, multiple obstacles have so far limited the benefit of CAR T cell therapy for patients with solid tumors. Novel manufacturing and engineering approaches show great promise to enhance CAR T cell function against solid tumors. However, similar to single agent chemotherapy approaches, CAR T cell monotherapy may be unable to achieve high cure rates for patients with difficult to treat solid tumors. Thus, combinatorial drug plus CAR T cell approaches may ultimately be required to achieve widespread clinical success. In this regard, we developed a novel high-content and high-throughput screen to evaluate 1114 FDA approved drugs to increase expression of the solid tumor antigen B7-H3 in metastatic osteosarcoma cells. In this proof-of-principle screen, we demonstrate that ingenol-3-angelate increased B7-H3 (CD276) mRNA, total protein, and cell surface expression. Mechanistically, ingenol-3-angelate increased B7-H3 expression via protein kinase C alpha activation. Functionally, ingenol-3-angelate induced B7-H3 expression enhanced B7-H3-CAR T cell function, highlighting utility of the approach, and paving the way for expanding this high-throughput and high-content technique to study other tumor and CAR T cell combinations.

## INTRODUCTION

The potential benefit of CAR T cell therapy for patients with cancer is most clearly demonstrated by the remarkable success of targeting CD19- and BCMA-positive malignancies.^1,2^ Despite this transformative progress, achieving similar benefits for patients with solid tumors has so far been limited.^3,4^ To enhance CAR T cell function against solid tumors, multiple obstacles need to be overcome including a lack of ideal target antigens, tumor heterogeneity, T cell trafficking to tumors, and immunosuppressive cells and molecules within the tumor environment.^5-7^

B7-H3 is an attractive immunotherapy target given its expression on multiple solid tumors with limited or no expression on most normal tissues.^8-12^ B7-H3-CAR T cells have potent antitumor activity without reported toxicities in multiple pre-clinical models.^8-10,13^ Results from these and other studies led to active B7-H3-CAR T cell therapeutic trials for patients with solid tumors (e.g. NCT04483778, NCT04670068),^14^ including a recently opened phase I study at St. Jude Children’s Research Hospital (NCT04897321).

Given the obstacles to success for CAR T cells against solid tumors, superior results may ultimately be obtained by developing rational combinatorial drug plus CAR T cell approaches. While some studies have evaluated drug plus CAR T cell combinations,^15-17^ novel high-throughput techniques capable of screening large numbers of compounds will likely enhance our ability to identify promising combinatorial treatment approaches for different solid tumor types.

Given these findings, we explored a novel, proof-of-principle, high-throughput, high-content screen to identify drugs with the ability to increase B7-H3 expression in metastatic osteosarcoma cells, which could potentiate B7-H3-CAR T cell effector function.

## RESULTS

### A novel high-content screen identifies an FDA approved drug to increase B7-H3 surface expression on metastatic osteosarcoma cells

To evaluate a proof-of-principle screen for identifying drugs with the potential to increase B7-H3 expression on metastatic osteosarcoma cells (LM7), we developed an immunofluorescence-based high-content assay to measure endogenous B7-H3 protein expression on the surface of single cells. B7-H3-positive LM7 cells^8^ were first seeded in 384-well optically-clear-bottom plates. Cells were then fixed but not permeabilized, stained with a B7-H3 antibody that recognizes an extracellular epitope, and imaged using a confocal-microscopy-based high-content screening system (CV8000) (**Fig. 1a**). To evaluate specificity, LM7 cells knocked out for B7-H3 (LM7-KO)^8^ served as a negative control. As expected, fluorescent signal was detected in LM7 but not LM7-KO cells, confirming specificity of the assay (**Fig. 1b**).

**Fig. 1.**
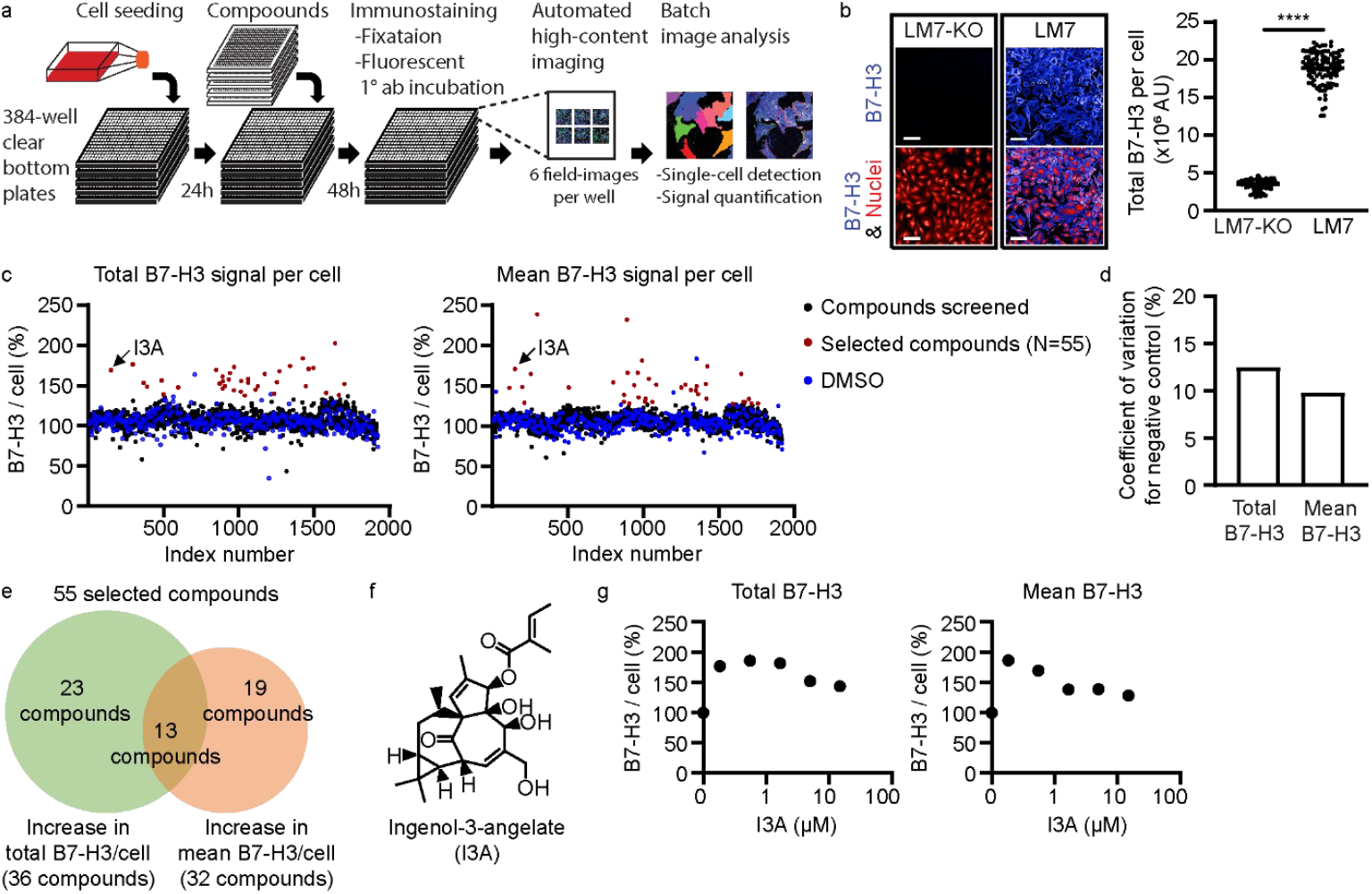
A high-content screen to identify FDA approved drugs that increase B7-H3 expression in metastatic osteosarcoma (LM7) cells. **a** Schematic of the high-content screen to quantify B7-H3 protein expression in a 384-well format. **b** Representative immunofluorescence images of B7-H3-negative (LM7-KO) and B7-H3-positive (LM7) cells (left panel; scale bars = 100μm) and quantification of total B7-H3 signal per cell (right panel). Each dot represents the average result of cells in a well (N=192 wells). **c** Total (left panel) and mean (right panel) B7-H3 expression per cell after exposure to 1114 FDA approved drugs. Results were normalized by averages of DMSO-treated wells in corresponding plates. Each dot represents the average result of cells in a well. **d** Coefficients of variation for negative controls of individual assay plates for total and mean B7-H3 signal per cell. **e** Venn diagram of drugs that increased total, mean, or both total and mean B7-H3 expression per cell. **f** The chemical structure of I3A. **g** The total and mean B7-H3 signal per cell after treatment with increasing doses of I3A. LM7 cells were treated with five 3-fold dilutions of 15μM I3A for 48 hours. Results are normalized by averages of DMSO-treated wells. Data represents mean ± SD. ****p<0.0001 by unpaired t-test (b). AU, arbitrary units.

After confirming specificity, we screened 1114 United States Food and Drug Administration (FDA) approved drugs to identify compounds that increase B7-H3 surface expression. LM7 cells were treated with each drug at a 10 μM concentration for 48 hours. Using the Columbus image analysis program, a batch analysis segmented single cells and measured B7-H3 fluorescent signals in each cell. Because some compounds alter cell size, which can affect output values, both the total and mean signal intensities per cell were determined to select compounds for in depth analysis (**Fig. 1c**). Because no compounds were known to increase cell-surface B7-H3 expression within 48 hours, the screen was processed without a positive control compound, and the total and mean B7-H3 signal per cell were normalized by a negative control (DMSO only). The average coefficients of variation for a negative control in total and mean B7-H3 signal per cell were 12.5% and 9.8% (**Fig. 1d**), indicating low measurement variation of the screening assay.

Using this method, 55 compounds were identified to increase the total and/or mean B7-H3 signal per cell (**Fig. 1c**). Of these 55 compounds, 23 increased the total signal per cell, 19 increased the mean signal per cell, and 13 increased both the total and mean B7-H3 signal per cell (**Fig. 1e**). After identifying these hits, a 5-point dose-response was measured at 24 and 48 hours post-treatment with each of these 55 compounds (**Supplementary Fig. 1a-d**), and 38 increased the total or mean B7-H3 signal per cell by greater than 50% for at least one concentration tested (**Supplementary Table 1**). One compound, ingenol-3-angelate (I3A; **Fig. 1f**) increased the total and mean B7-H3 signal per cell by up to 100% at 48 hours (**Fig. 1g**). I3A induced maximal B7-H3 expression at 0.19-1.67 μM, with lower induction above these concentrations, demonstrating a biphasic response.

### I3A induced B7-H3 expression occurs at both the mRNA and protein levels

To further evaluate the effect of I3A treatment on B7-H3 expression, LM7 cells were treated with 22 doses of I3A for 6, 24, and 48 hours. Consistent with the 5-dose response experiment, B7-H3 surface expression increased in a biphasic manner when treated for 24 or 48 hours, whereas treatment for 6 hours had limited effect (**Fig. 2a**). I3A concentrations of 0.1 to 0.5 μM increased B7-H3 expression, while higher concentrations had less of an effect (**Fig. 2a**). Additionally, I3A is not an autofluorescent molecule that interferes with the immunofluorescent assay because the signal intensity of fluorophore-conjugated isotype control IgG was minimal with I3A treatment (**Fig. 2a**).

**Fig. 2.**
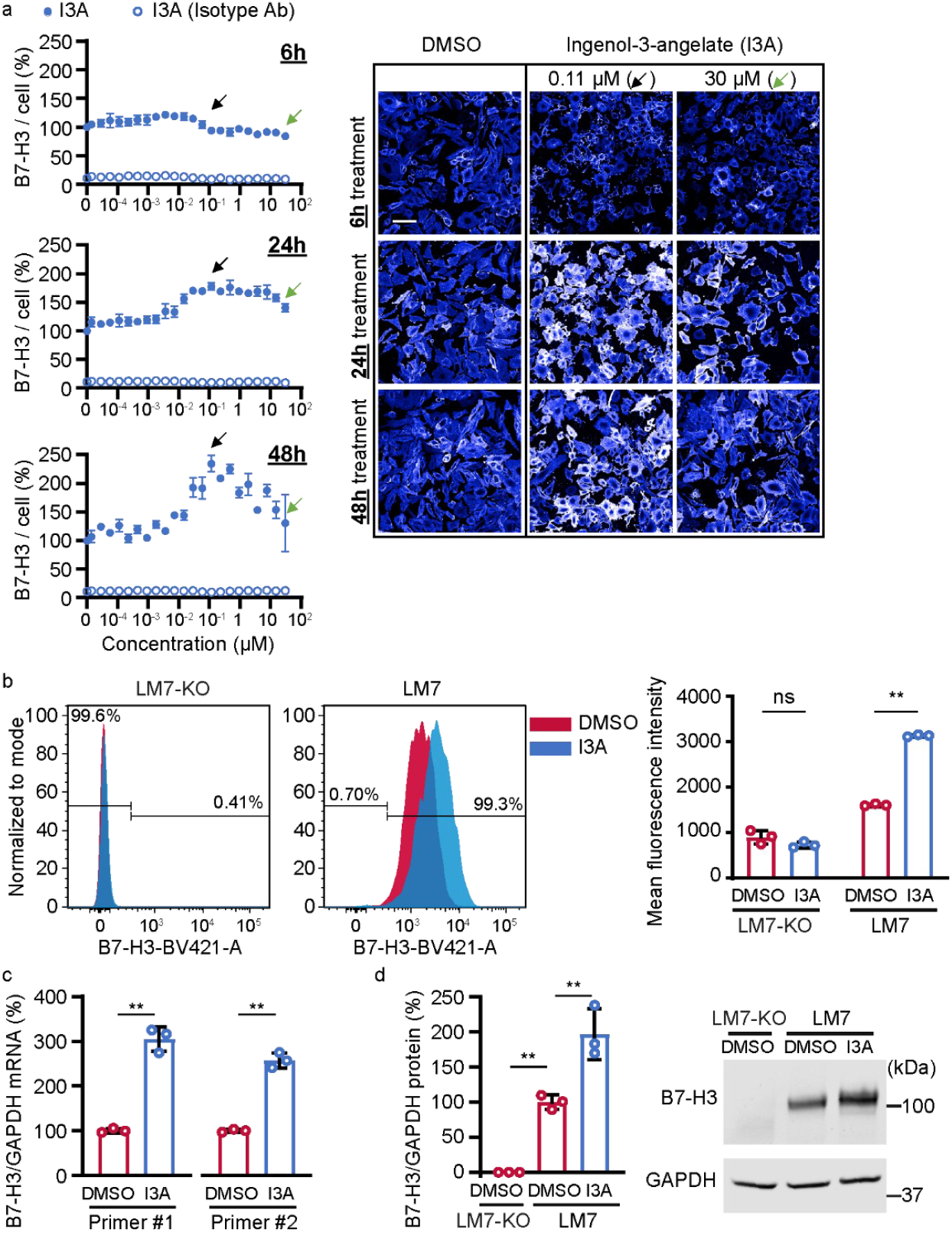
I3A increases B7-H3 mRNA, total protein, and cell-surface protein expression in LM7 cells. **a** B7-H3 expression per cell after treatment with 22 doses of I3A for 6, 24, or 48 hours. Total B7-H3 signal per cell (left panel) was normalized by averages of DMSO-treated wells. Filled circles represent detection by B7-H3-specific antibody and unfilled circles isotype IgG control. Black arrows represent treatment with 0.11 μM I3A, and green arrows represent treatment with 30 μM I3A (N=4). Representative LM7 fluorescent images (right panel) after treatment with indicated concentrations of I3A at 6, 24, or 48 hours are shown (scale bar = 100 μm). **b** Flow cytometry evaluation of B7-H3 surface expression on LM7-KO or LM7 cells treated with DMSO or 0.1 μM I3A. Representative flow plots (left panel) and summary of replicates (right panel; N=3) are shown. **c** B7-H3 mRNA expression in LM7 cells post-treatment with DMSO or 0.5 μM I3A for 48 hours (N=3). B7-H3 mRNA expression was measured by RT-qPCR with two different primers and normalized by the GAPDH mRNA expression and normalized again by averages of DMSO-treated samples. **d**. B7-H3 protein in LM7 cells after treatment with DMSO or 0.5 μM I3A for 48 hours (N=3). Relative B7-H3 protein expression to GAPDH protein expression was normalized by DMSO-treated LM7 samples. Summary data (left panel) and representative western blot (right panel) are shown. Data represents mean ± SD. **p<0.01, ns, non-significant by two-way ANOVA (b), unpaired t-test (c), or one-way ANOVA (d).

To validate these findings in a format used for testing CAR T cell function, LM7 or LM7-KO cells were treated with 0.1 μM I3A or DMSO control for 48 hours in 24 well plates, and B7-H3 surface expression was measured by flow cytometry. Whereas no B7-H3 was detected in DMSO or I3A treated LM7-KO cells, I3A significantly increased B7-H3 surface expression on LM7 cells in this setting, validating our high-content screening results (**Fig. 2b**).

We next queried if increased B7-H3 expression occurs at the mRNA level, protein level, or both. RT-qPCR using two predesigned commercial TaqMan probes showed that 0.5 μM I3A increased B7-H3 mRNA expression at 48 hours (**Fig. 2c**). Western blot analysis showed that total B7-H3 protein expression was also increased by 0.5 μM I3A at 48 hours (**Fig. 2d**). The B7-H3 protein bands were specific because LM7-KO cell lysates did not show a B7-H3 band. Intriguingly, I3A induced upregulation of B7-H3 expression was cell type-specific. When A549 (lung adenocarcinoma) cells were treated with I3A, no increase in B7-H3 expression was detected at 24 or 48 hours in a 12-dose immunofluorescence assay (**Supplementary Fig. 2a**). In addition, RT-qPCR demonstrated that 0.5 μM I3A did not change B7-H3 mRNA expression in A549 cells (**Supplementary Fig. 2b**). These results demonstrate that I3A increased B7-H3 expression at the mRNA and protein level in a cell type-specific manner.

### I3A pre-treatment enhances B7-H3-CAR T cell function against metastatic osteosarcoma cells

We next asked if pre-treatment of LM7 cells with I3A enhances B7-H3-CAR T cell effector function. To evaluate this, LM7 cells were pre-treated with 0.1 μM I3A or DMSO control for 48 hours, then washed and plated in a 24 well plate for 4 hours to adhere, followed by addition of B7-H3-CAR T cells. CAR T cells specific for CD19 served as negative controls (Control-CAR). While limited cytokine secretion was detected for both Control-or B7-H3-CAR T cells against LM7-KO, I3A treatment of LM7 cells significantly enhanced B7-H3-CAR T cell IFNγ and IL-2 secretion (**Fig. 3a**). To evaluate cytotoxicity, LM7-KO or LM7 cells were pre-treated with 0.1 μM of I3A and tumor cell killing was quantified using an impedance-based assay (xCelligence). Given that higher CAR T cell doses may efficiently kill tumor cells despite lower antigen expression, we evaluated cytotoxicity in T cell dose-titration assays (**Fig. 3b**). While I3A enhanced B7-H3-CAR T cell cytotoxicity at all T cell-to-target cell ratios evaluated, the difference was most pronounced at the lowest ratio tested (1 T cell to 8 tumor cells; **Fig. 3c**). Taken together, these results confirm that I3A induced B7-H3 expression improves B7-H3-CAR T cell effector function against metastatic osteosarcoma cells.

**Fig. 3.**
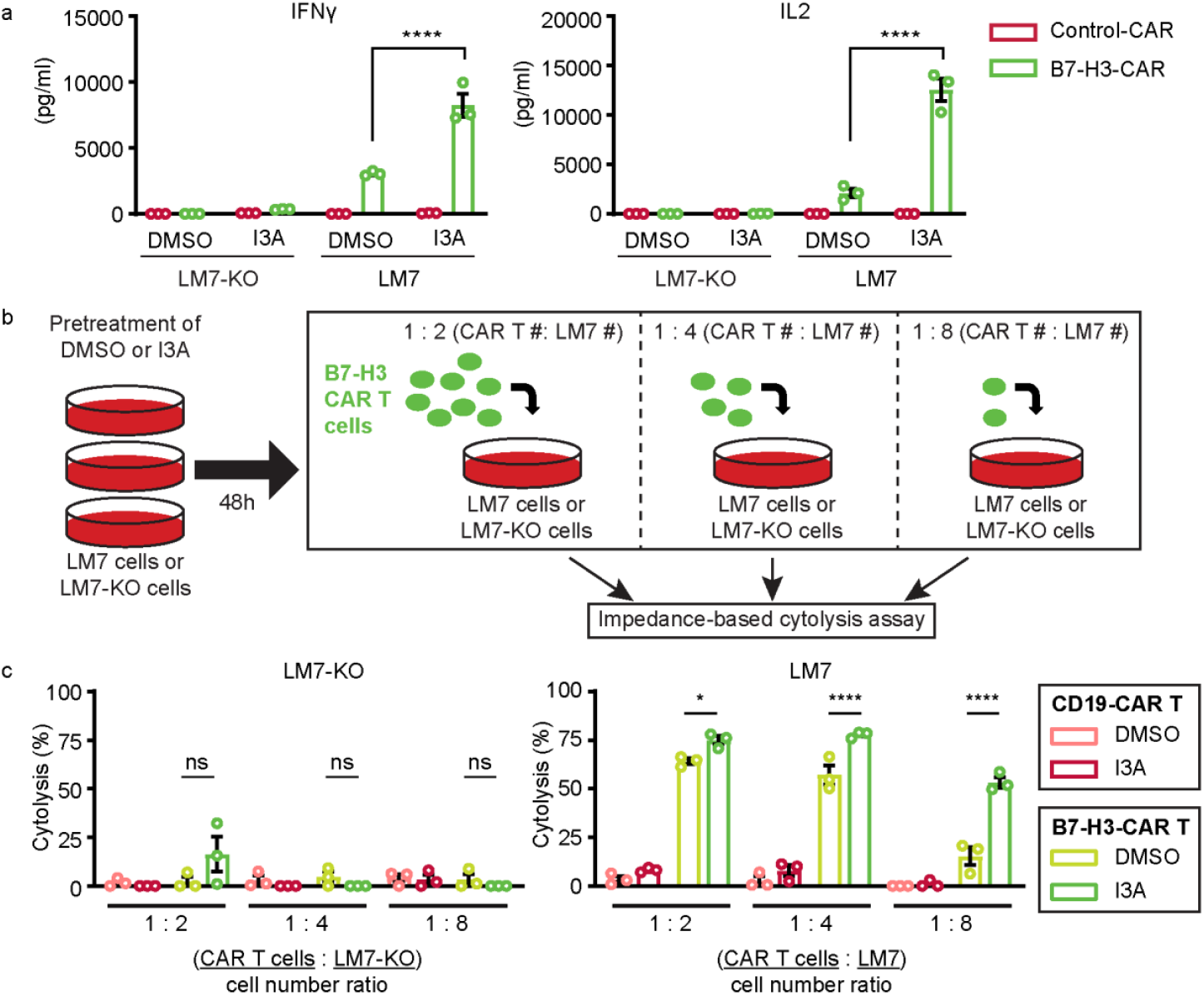
I3A induced B7-H3 expression enhances B7-H3-CAR T cell effector function. **a** CD19-CAR (Control) or B7-H3-CAR T cell IFNγ (left panel) and IL2 (right panel) secretion post coculture with LM7-KO or LM7 cells treated with DMSO or 0.1 μM I3A (N=3). **b** Diagram of cytotoxicity assay. **c** Control- or B7-H3-CAR killing of LM7-KO or LM7 cells treated with DMSO or 0.1μM I3A (N=3). Data represents mean ± SEM (a and c). *p<0.05, ****p<0.0001, ns, non-significant by two-way ANOVA (a and c).

### I3A increases B7-H3 expression via protein kinase C (PKC) activation

We next investigated the mechanism behind I3A-induced B7-H3 expression. I3A is a PKC agonist,^18^ raising the possibility that I3A increases B7-H3 expression by modulating PKC activity. Supporting this hypothesis, phorbol 12,13-dibutyrate (PDBu), another PKC activator, increased B7-H3 protein expression in a biphasic manner similar to I3A in the immunofluorescence assay (**Fig. 4a**).

**Fig. 4.**
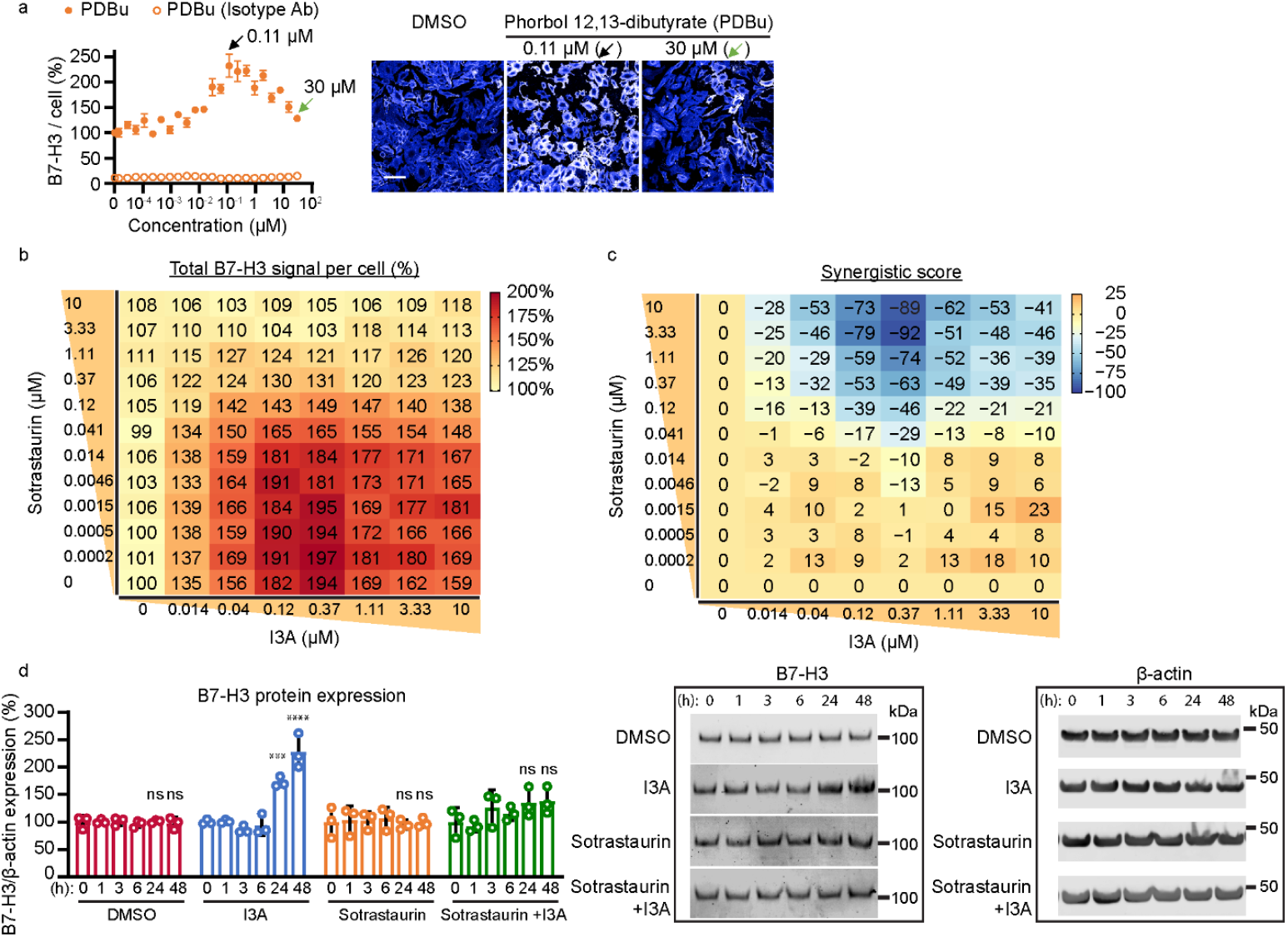
PKC inhibition abrogates I3A induced B7-H3 protein expression. **a** LM7 cells were treated with 22 doses of PDBu and evaluated with the immunofluorescence screen. Total B7-H3 signal per cell (left panel; N=4) was normalized by averages of DMSO-treated wells. Filled circles represent detection by B7-H3-specific antibody and unfilled circles isotype IgG control. Representative LM7 fluorescent images (right panel) after treatment with indicated concentrations of PDBu are shown. Immunofluorescence analysis for the combinational treatment of 8 doses of I3A and 12 doses of sotrastaurin was performed. **b** B7-H3 signal heatmap and **c** synergistic score heatmap (highest single agent model) are shown. The total B7-H3 signal per cell was normalized by DMSO-treated samples. Data are shown as means of quadruplicates. Negative synergistic scores indicate antagonistic effects. **d** Western blot analysis for the combinational treatment of I3A and sotrastaurin. B7-H3 protein expression relative to β-actin normalized by 0 hour treated samples (left panel) and representative western blots (middle and right panels) are shown. Data represents mean ± SD (a and d). ***p<0.001, ****p<0.0001, ns, non-significant by one-way ANOVA (d).

To further evaluate the role of PKC in I3A induced B7-H3 expression, we asked if sotrastaurin, a pan-PKC inhibitor, abrogates increased B7-H3 expression due to I3A. To test this, LM7 cells were treated with 96 combinations of sotrastaurin (12 doses) and I3A (8 doses) for 48 hours, and the total B7-H3 signal per cell was measured using the immunofluorescence assay. The average of quadruplicated results was normalized by the basal fluorescence levels in DMSO-treated samples. As expected, I3A alone increased B7-H3 expression in a biphasic manner, whereas sotrastaurin alone had no effect (**Fig. 4b, Supplementary Fig. 3**). Importantly, when used in combination, sotrastaurin suppressed I3A-induced B7-H3 expression in a dose-dependent manner, and almost completely abolished I3A induced B7-H3 expression at concentrations of 3.33 μM and above (**Fig. 4b**). To quantify the antagonistic effect of sotrastaurin on I3A, the highest-single agent (HSA) model^19^ was applied to the 96-combinational treatment results. Negative HSA model scores, at a range of 0.04-10 μM I3A and ≥ 0.12 μM sotrastaurin, represent the antagonistic effect of I3A and sotrastaurin (**Fig. 4c**).

To further evaluate these findings, we implemented time-course western blot experiments using combinatorial I3A and sotrastaurin treatment. LM7 cells were treated with DMSO, 0.5 μM I3A, or 5 μM sotrastaurin only, or a combination of 0.5 μM I3A and 5 μM sotrastaurin for different lengths of time. Western blot results confirmed the PKC inhibitor sotrastaurin abolished I3A induced B7-H3 expression at 24 and 48 hours post-treatment (**Fig. 4d**). These results demonstrate that I3A induces B7-H3 expression in LM7 cells via PKC activation.

### Knockdown of PKCα abrogates I3A induced B7-H3 expression

Since there are 9 PKC isoforms (α, β, γ, δ, ε, ζ, η, θ, and ι), we asked if one or more isoforms are responsible for I3A induced B7-H3 expression. First, mRNA expression of each isoform was measured in LM7 cells by RT-qPCR. Transcripts for PKCα were highly expressed compared to the other 8 isoforms (**Fig. 5a**). Intriguingly, while I3A treatment did not change the PKCα mRNA level in LM7 cells (**Supplementary Fig. 4**), it reduced PKCα protein expression by 90% after 48 hours, as determined by western blot analysis (**Fig. 5b**). These results are consistent with previous reports that PKC activators reduce PKC protein levels by enhancing PKC degradation.^18,20,21^

**Fig. 5.**
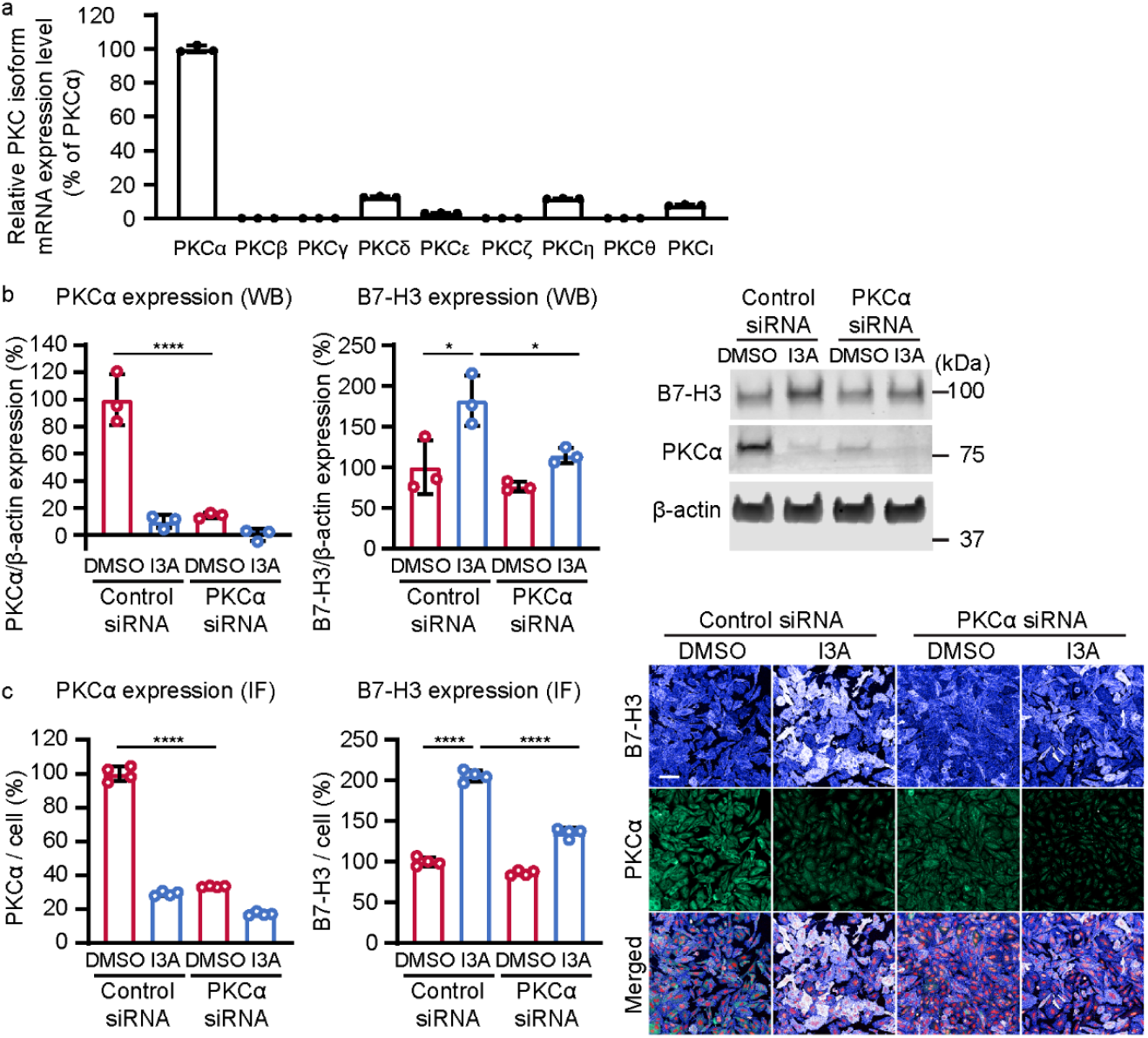
PKCα is required for I3A induced B7-H3 protein expression. **a** mRNA expression of 9 PKC isoforms measured by RT-qPCR (N=3). The mRNA expression levels were normalized by the PKCα mRNA level. After knockdown with control or PKCα siRNA, B7-H3 and PKCα protein levels were determined by western blot and immunofluorescence analysis. **b** Summary B7-H3 and PKCα protein expression determined by western blot after siRNA (left and middle panels; N=3) and representative western blot (right panel) are shown. **c** Summary B7-H3 and PKCα protein expression determined by immunofluorescence assay after siRNA (left and middle panels; N=4) and representative images (right panel) are shown. B7-H3 (extended blue), PKCα (green), and nuclei (red). Scale bar = 100μm. Data represents mean ± SD (a, b, and c). *p<0.05, ****p<0.0001 by two-way ANOVA (b and c).

To determine if PKCα plays a direct role in I3A induced B7-H3 expression, we knocked down PKCα in LM7 cells by transfecting a pool of 4 siRNAs targeting PKCα. siRNA knockdown successfully reduced PKCα protein expression by 85% determined by western blot (**Fig. 5b**) and 67% by immunofluorescence assay (**Fig. 5c**). As expected, I3A treatment in the presence of non-targeting control siRNA increased B7-H3 protein expression by 82% by western blot (**Fig. 5b**) and 100% by immunofluorescence assay (**Fig. 5c**). Strikingly, PKCα knockdown abrogated I3A induced B7-H3 expression (**Fig. 5b**,**c**). In the setting of PKCα knockdown, I3A increased B7-H3 expression by only 38% in western blot (**Fig. 5b**) and 50% in immunofluorescence assays (**Fig. 5c**), demonstrating that I3A increased B7-H3 expression in LM7 cells via PKCα.

## DISCUSSION

Here we describe a rational approach to screen FDA approved drugs for upregulating tumor target antigen expression to enhance CAR T cell activity. Using this approach, I3A was identified to upregulate B7-H3 in a cell type-specific manner, which occurred at both the mRNA and protein levels. Mechanistically, I3A upregulated B7-H3 through PKC activation, and this effect was diminished specifically by PKCα knockdown. Importantly, I3A induced B7-H3 expression enhanced B7-H3-CAR T cell effector function against metastatic osteosarcoma cells. Thus, our findings demonstrate utility of this proof-of-principle screen to identify FDA approved drugs to upregulate target antigen expression on cancer cells to enhance CAR T cell activity.

Large scale screens will likely aid the discovery of rational combinations to improve response rates for patients treated with CAR T cells. We implemented a screen using over 1000 FDA approved drugs to modulate surface antigen expression in metastatic osteosarcoma cells. Our confocal-microscopy-based high-content screen measured endogenous B7-H3 expression in unmodified cells in a 384-well plate format, which can be used to screen a large number of compounds with various types of cells in a high-throughput manner. While limited in number, other important studies have begun to shed light on the promise of large-scale screens to identify combinatorial approaches. In a screen using over 500 compounds, SMAC mimetics enhanced CD19-CAR T cell activity against B-cell acute lymphoblastic leukemia (B-ALL) and diffuse large B cell lymphoma. SMAC mimetics did not increase CD19 expression in primary B-ALL cells, but enhanced CAR T cell killing by modulating death receptor mediated apoptosis pathways. Similar to our findings, the SMAC mimetic/CD19-CAR T cell combinatorial benefit was cell line dependent.^22^ Other examples of drug plus CAR T cell screens have identified hedgehog pathway inhibitors (e.g. JK184) to enhance B7-H3-CAR T cell killing of breast cancer cells via inhibiting tumor resistance to apoptosis,^23^ and insulin-like growth factor receptor 1/insulin receptor inhibitors (BMS-754807, linsitinib) to enhance GD2-CAR T cell activity against diffuse midline glioma.^24^ Overall, there is a growing body of evidence demonstrating that cutting edge technologies can inform rational combinatorial CAR T cell treatment strategies for multiple cancer types.

Aside from large scale screens, individual combinatorial drug plus CAR T cell approaches have been evaluated, and some translated into clinical trials.^15-17,25^ We identified I3A as a drug to increase B7-H3 expression in metastatic osteosarcoma cells. While this demonstrated utility of our screen, I3A is a topical medication and not readily applicable for patients with osteosarcoma. Multiple CAR T cell products have been combined with checkpoint blockade therapy (e.g. PD-1 antibody) with resultant enhanced anti-solid tumor activity.^26,27^ Active or recently completed clinical trials are evaluating this approach for patients with solid tumors (e.g. NCT04995003, NCT03726515).^28^ Other agents including radiation therapy^29^, lenalidomide,^30^ and hypomethylating agents such as decitabine have confirmed benefit in pre-clinical studies and are being evaluated in combination with CAR T cells in the clinic (e.g. NCT03017131). Intriguingly, the HDAC inhibitor vorinostat (SAHA) increases B7-H3 expression in solid tumor cells approximately 5 days after treatment, leading to enhanced B7-H3-CAR T cell activity.^31^ In our screen, vorinostat was not detected to upregulate B7-H3 in LM7 cells, likely because we evaluated agents to upregulate B7-H3 within 48 hours after treatment. In total, multiple combinatorial approaches offer promise to enhance the antitumor activity of CAR T cells against difficult to treat cancers and are likely to inform the next generation of CAR T cell clinical trials for patients with solid tumors.

While promising, combinatorial drug plus CAR T cell approaches pose challenges. Safety considerations must be considered when developing combinatorial treatment strategies. The goal of this work was to develop a proof-of-principle screening strategy to identify FDA approved drugs to increase B7-H3 expression, so normal cell lines were not included in our initial experiments. For subsequent studies, identifying representative normal cells or testing relevant in vivo models will be critical to establish specificity and safety. Timing of drug delivery in relation to CAR T cell infusion is another important challenge. Given that some promising combinatorial agents may enhance T cell activity and others impair T cell function, rational treatment regimens will need to be carefully designed to maximize the potential impact of such approaches.

In conclusion, high-throughput, high-content screening is a promising strategy to identify compounds and mechanisms to enhance the efficacy of CAR T cell therapies for patients with solid tumors. Our novel large-scale image-based screening approach (high-throughput and high-content) is an effective tool to identify agents to upregulate target antigen expression on tumor cells and adds to the growing body of evidence that combinatorial treatment strategies can be developed and successfully implemented to move the field forward.

## MATERIALS AND METHODS

### Cell culture and compounds

The metastatic osteosarcoma cell line, LM7, was kindly provided by Dr. Eugenie Kleinerman (MD Anderson Cancer Center, Houston, TX). The A549, (lung cancer) cell line was purchased from ATCC. Tumor cells were cultured with conventional cell culture methods. DMEM (Cytiva, SH30022.01) with GlutaMAX (Gibco, 51985-034) and 10% FBS (Cytiva, SH300700.03TIR) was used for the cell culture. All cells were maintained at 37°C in 5% CO_2_. Cell lines were authenticated by short tandem repeat profiling using the service of the ATCC (FTA sample collection kit) and routinely checked for mycoplasma using the MycoAlert mycoplasma detection kit (Lonza). Ingenol-3-angelate (I3A) (Cayman, 16207), phorbol 12,13-dibutylate (PDBu) (Torcris, 4153), sotrastaurin (MCE, HY-10343/CS0090) were treated as described.

### B7-H3 knockout cells

B7-H3 knockout LM7 cells were generated as previously described.^8^ B7-H3 knockout A549 cells (A549-KO) were generated using CRISPR-Cas9 technology. Briefly, 400,000 A549 cells were transiently transfected with precomplexed ribonuclear proteins (RNPs) consisting of 150pmol of chemically modified sgRNA (5’ – UCUCCAGCACACGAAAGCCA - 3’, Synthego), 35pmol of Cas9 protein (St. Jude Protein Production Core), and 500ng of pMaxGFP (Lonza) via nucleofection (Lonza, 4D-Nucleofector™ X-unit) using solution P3 and program CM-130 in a small (20ul) cuvette according to the manufacturer’s recommended protocol. Five days post nucleofection, cells were single-cell sorted by FACs to enrich for GFP+ (transfected) cells, clonally selected, and verified for the desired targeted modification via targeted deep sequencing using gene specific primers with partial Illumina adapter overhangs (hB7-H3.F – 5’ ATTCATAGTGTTAGGGCCCAGTGAGG -3’ and hB7-H3.R – 5’ GCCCCATCTGCACACACACACTCGT -3’, overhangs not shown) as previously described.^32^ Briefly, cell pellets of approximately 10,000 cells were lysed and used to generate gene specific amplicons with partial Illumina adapters in PCR#1. Amplicons were indexed in PCR#2 and pooled with targeted amplicons from other loci to create sequence diversity. Additionally, 10% PhiX Sequencing Control V3 (Illumina) was added to the pooled amplicon library prior to running the sample on an Miseq Sequencer System (Illumina) to generate paired 2 × 250bp reads. Samples were demultiplexed using the index sequences, fastq files were generated, and NGS analysis of clones was performed using CRIS.py.^32^ Two hB7-H3 knockout clones containing only out-of-frame indels were identified. Final clones were authenticated using the PowerPlex® Fusion System (Promega) performed at the Hartwell Center (St. Jude), and one clone chosen for further use. This A549-KO clone was expanded, stained with B7-H3 antibody (clone 7-517; Becton Dickinson, Franklin Lakes, NJ, USA) and again single-cell sorted on the B7-H3-negative population using a BD FACSAriaIII instrument to ensure a pure B7-H3 negative population. After producing the final A549-KO product, cells were confirmed B7-H3 negative by flow cytometry, authenticated by short tandem repeat profiling using the service of the American Type Tissue Collection (FTA sample collection kit; ATCC, Manassas, VA, USA), and tested negative for mycoplasma by the MycoAlert™Plus Mycoplasma Detection Kit (Lonza).

### High-content imaging screening assay

LM7 cells were seeded in 384-well plates with an optically clear bottom (PerkinElmer, PDL-coated CellCarrier-384 Ultra). The next day, 1114 compounds in an FDA-approved drug library (St. Jude Children’s Research Hospital) (∼10 μM) were dispensed to the plates of cells using an Echo 555 Liquid Handler (Labcyte). After 48 hours in a cell culture incubator, cells were washed with PBS three times using ELx405 plate washer (BioTek). Four % paraformaldehyde (Electron Microscopy Sciences, 15710) in PBS was incubated for 15 minutes at room temperature. Cells were washed with PBS three times. 1:2400 BV421 anti-B7-H3 antibody (BD Biosciences, 565829), 1:10,000 DRAQ5 (Thermo Scientific, 62251), 1:1000 WGA Alexa Fluor 488 (Invitrogen, W11261), and 2% horse serum (Gibco, 26050070) in PBS was incubated overnight at 4°C. Cells were washed with PBS three times.

Using Max, an automated robot system in St. Jude Children’s Research Hospital, immunostained cells in the microwell plates were imaged by CV8000, a confocal microscopy-based high-content screening system (Yokogawa). Images were imported to Columbus, an imaging analysis software (PerkinElmer). Using Columbus, each cell in the images was segmented using DRAQ5 and/or WGA signal, and the mean and total signal intensity of B7-H3 per cell was measured. Coefficients of variation were calculated using Genedata Screener software (Genedata).

### Immunofluorescence imaging assay

Immunofluorescence for B7-H3 was performed as described above in the high-content imaging screening assay method section. For immunostaining PKCα, cells were permeabilized by 0.1% TritonX-100 for 15 minutes after fixation. 1:100 anti-PKCα antibody (Cell Signaling Technology, 2056S) 1:10,000 DRAQ5 (Thermo Scientific, 62251), 1:1000 WGA Alexa Fluor 488 (Invitrogen, W11261), and 2% horse serum (Gibco, 26050070) in PBS was incubated overnight at 4°C. After washing with PBS 3 times, 1:500 Alexa Fluor Plus 555 anti-rabbit IgG antibody (Invitrogen, A-31572) was incubated for 1 hour.

### Flow cytometry

A FACSCanto II (BD) instrument was used to acquire flow cytometry data, which was analyzed using FlowJo v10 (BD). For surface staining, samples were washed with and stained in PBS (Lonza) with 1% FBS (GE Healthcare Life Sciences). For all experiments, matched isotypes or known negatives (e.g. non-transduced T cells) served as gating controls. CAR detection was performed using F(ab’)_2_ fragment specific antibody (polyclonal, Jackson ImmunoResearch, West Grove, PA). Tumor cell lines were evaluated for expression of B7-H3 using B7-H3 antibody (clone 7-517, BD or clone FM276, Miltenyi).

### Analysis of cytokine production

Tumor cells were pre-treated with 0.1 μM I3A or DMSO control for 48 hours, then washed and plated in a 24 well plate for 4 hours to adhere, followed by addition of T cells. 2.5×10^5^ T cells were added to 5×10^5^ LM7 or LM7-KO treated with I3A or DMSO. Approximately 24 hours post-coculture, supernatant was collected and frozen for subsequent analysis. IFNγ and IL-2 production were measured using a quantitative ELISA per the manufacturer’s instructions (R&D Systems, Minneapolis, MN).

### In vitro cytotoxicity assay

The xCELLigence RTCA MP instrument (Agilent, Santa Clara, CA) was used to assess CAR T cell cytotoxicity. All assays were performed in triplicate without the addition of exogenous cytokines. First, 30,000 LM7 or LM7-KO cells were added to each well of a 96 well E-Plate (Agilent) in complete RPMI, and allowed to adhere overnight to reach a cell index (relative cell impedance) plateau. Next, complete RPMI plus 0.1 μM I3A or DMSO was added for 48 hours. Thereafter, I3A and DMSO were removed, and CAR T cells in complete RPMI added at 15,000, 7500, or 3750 cells per well. Cell index was monitored every 15 minutes for 3 days and normalized to the maximum cell index value immediately prior to T cell plating. Percent cytotoxicity was calculated using RTCA Software Pro immunotherapy module (Agilent).^33^

### Western blotting

Cells were seeded in 6-well plates. Cells were washed with cold PBS on ice. Cells were scraped with cold RIPA buffer (Pierce, 89901) and Halt protease and phosphatase inhibitor cocktail (Pierce, 78440) and vortexed. After 15-minute centrifugation at 16,000 g at 4°C, the supernatant was kept, and total protein concentration was measured by BCA Protein Assay Kit (Pierce, 23227) following the manufacture’s instruction. Protein samples for western blot assay were prepared with the same total protein amount and the sample volume in each experiment and boiled at 95°C for 5 minutes.

The protein samples were loaded and run on 4-12% Bis-Tris gels (Invitrogen). Proteins were transferred on nitrocellulose membranes using iBlot and iBlot2 system (Invitrogen). Transferred blots were blocked with Odyssey PBS blocking buffer (LI-COR, 927-40000) for one hour at room temperature. 1:1000 anti-B7-H3 antibody (Cell Signaling Technology, 14058S) or 1:1000 anti-PKC-alpha antibody (Cell Signaling Technology, 2056S) with or without 1:10,000 anti-beta actin (Sigma, A5441) or 1:1000 anti-GAPDH antibodies (Cell Signaling Technology, 97166S) in Odyssey PBS blocking buffer overnight at 4°C. After washing with TBST three times, membranes were incubated with 1:5000 IRDye 800CW anti-rabbit IgG (LI-COR, 926-32211) with or without 1:5000 IRDye 680RD anti-mouse IgG (LI-COR, 926-68070) in Odyssey PBS blocking buffer for one hour at room temperature. Blots were imaged using Odyssey CLx (LI-COR) and quantified using Image Studio software (LI-COR).

### RT-qPCR

Cells were seeded in 6-well plates. Cells were washed with PBS and incubated with Trypsin-EDTA at 37 °C until detached. Detached cells were neutralized with culture medium and centrifuged at 200 *g* for two minutes. The supernatant was removed. From the cell pellets, RNA was extracted using Maxwell simplyRNA (Promega). RNA concentration was measured by Nanodrop 8000 (ThermoScientific). cDNA was generated with 1 μg RNA using Superscript VILO (Invitrogen). qPCR was performed using Taqman Fast Advanced Master Mix (Applied Biosystems) with predesigned primers (Invitrogen, 4331182) (Primers’ Assay IDs: B7-H3 [Hs00987207_m1 and Hs00228846_m1]; GAPDH [Hs02786624_g1]; PKCα [Hs00925200_m1]; PKCβ [Hs00176998_m1]; PKCγ [Hs00177010_m1]; PKCδ [Hs01090047_m1], PKCε [Hs00942886_m1], PKCζ [Hs00177051_m1], PKCη [Hs00178933_m1]; PKCθ [Hs00234709_m1]; PKCι [Hs00995852_g1])

### Knockdown with siRNA

PKCα siRNA (ON-TARGETplus SMARTpool PRKCA [Dharmacon, L-003523-00-0005, 5578]) and non-targeting control siRNA (ON-TARGETplus Non-targeting Pool [Dharmacon, D-001810-10-05]) were used for the knockdown.

siRNAs were transfected by reverse-transfection using Lipofectamine RNAiMAX (Invitrogen, 100014474). 1.5 pmol (27.5 nL of 20 μM, 384-well plates) or 75 pmol (1.375 μL of 20 μM, 6-well plates) siRNA and 10 μL (384-well plates) or 500 μL (6-well plates) Lipofectamine RNAiMAX in OptiMEM (Gibco, 51985-034) were incubated for 20 min at room temperature. 15 μL (384-well plates) or 750 μL cells (6-well plates) were added to the siRNA and RNAiMAX mixture and incubated overnight in a cell incubator. 25 μL fresh medium was added to 384-well plates or medium was replaced with fresh medium in 6-well plates.

### Combinational matrix treatment for antagonistic effect

The next day after cell seeding in a 384-well plate (PerkinElmer, PDL-coated CellCarrier-384 Ultra), combinations of DMSO or 11 does of sotrastaurin and DMSO or 7 doses of ingenol-3-angelate were dispensed in quadruplicate using an Echo555 (Labcyte). Immunostaining and quantification of the B7-H3 protein expression level were performed as described above in the high-content imaging screening assay method section. Quadruplicated results were averaged and normalized and then analyzed for the activities and Highest Single Agent model scores using SynergyFinder R package ^34^.

### Statistical analysis

Statistical significance was calculated by GraphPad Prism (GraphPad Software). For comparison of two populations, statistical significance was determined by t-test with two sides. For comparison of more than two populations within a group, statistical significance was determined by one-way ANOVA. For comparison with multiple groups, statistical significance was determined by two-way ANOVA. A p-value cutoff of 0.05 was used to establish statistical significance.

## DATA AVAILABILITY

All data supporting the findings of this study are available from the corresponding authors upon reasonable request.

## ACKNOWLEDGEMENTS

We thank the Compound Management Center at the Department of Chemical Biology and Therapeutics for providing the FDA-approved drug library. We thank the Center for Advanced Genome Engineering (CAGE) at St. Jude for assisting to create knockout cell lines used in this work. The CAGE is supported by ALSAC and the NCI grant P30 CA021765. This work was supported by the Assisi Foundation of Memphis to CD, and ALSAC to TC and CD.

## CONFLICT OF INTERESTS

CD is a co-inventor on patent applications in the field of T cell therapy for cancer. All other authors declare no competing interests.

## CONTRIBUTIONS

Conceptualization: HWL, ANB, TC, CD. Formal Analysis: HWL, TC, CD. Funding acquisition: TC, CD. Investigation: HWL, CO, DGC. Methodology: HWL, CO, TC, CD. Project administration: TC, CD. Supervision: TC, CD. Validation: HWL, CO, TC, CD. Visualization: HWL, CO, TC, CD. Writing – original draft: HWL, CO, TC, CD. Writing – review & editing: HWL, CO, ANB, DGC, TC, CD.

