## Supplementary Information for "A high-content screen identified ingenol-3-angelate as an enhancer of B7-H3-CAR T cell activity by increasing B7-H3 protein expression on the target cell surface via PKCα activation"

**This PDF file includes:**

Figures. S1-S4
Table S1

**Figure. S1.**

a. Total B7-H3 signal / cell (%), 24h treatment

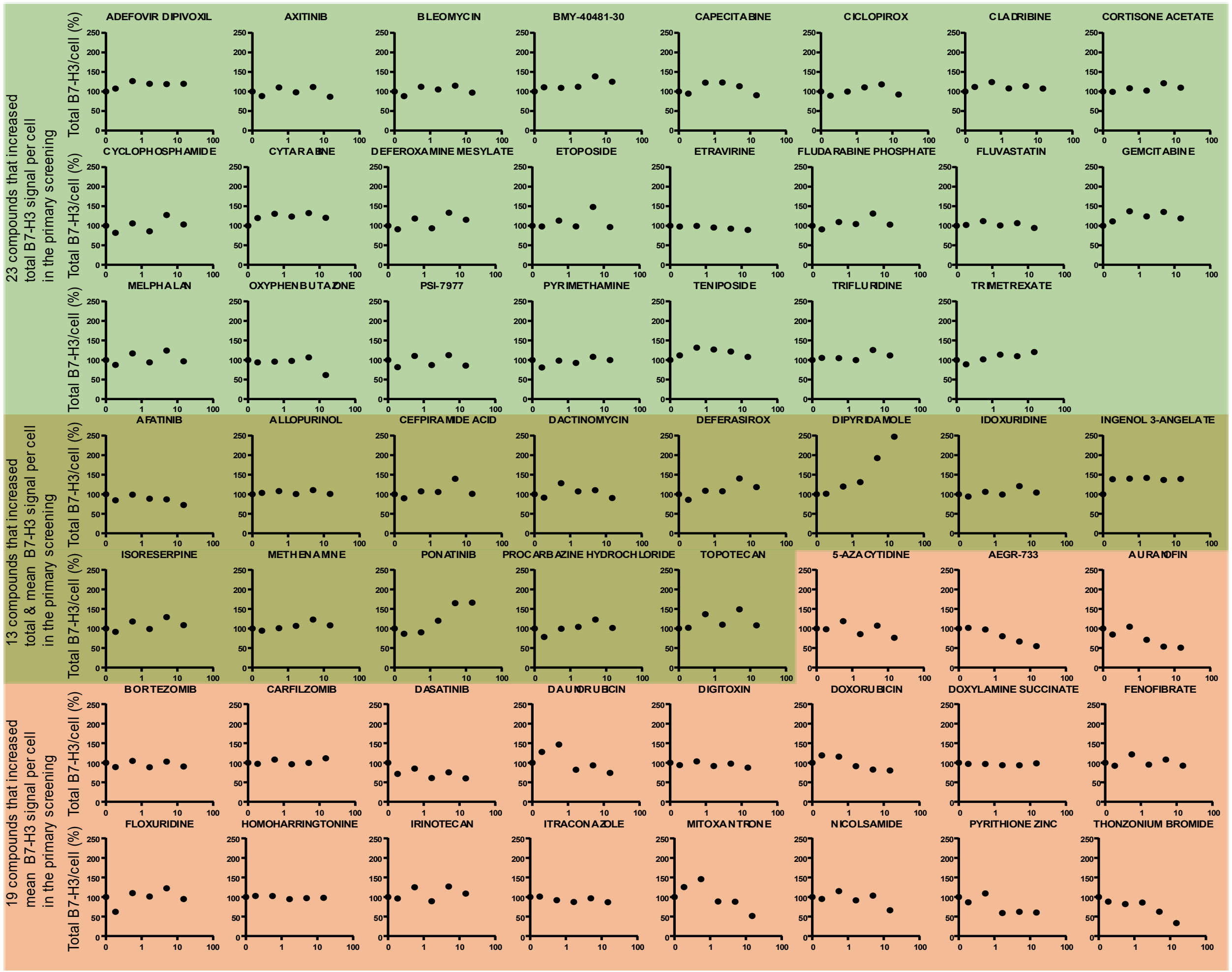

**Figure. S1.**

b. Mean B7-H3 signal / cell (%), 24h treatment

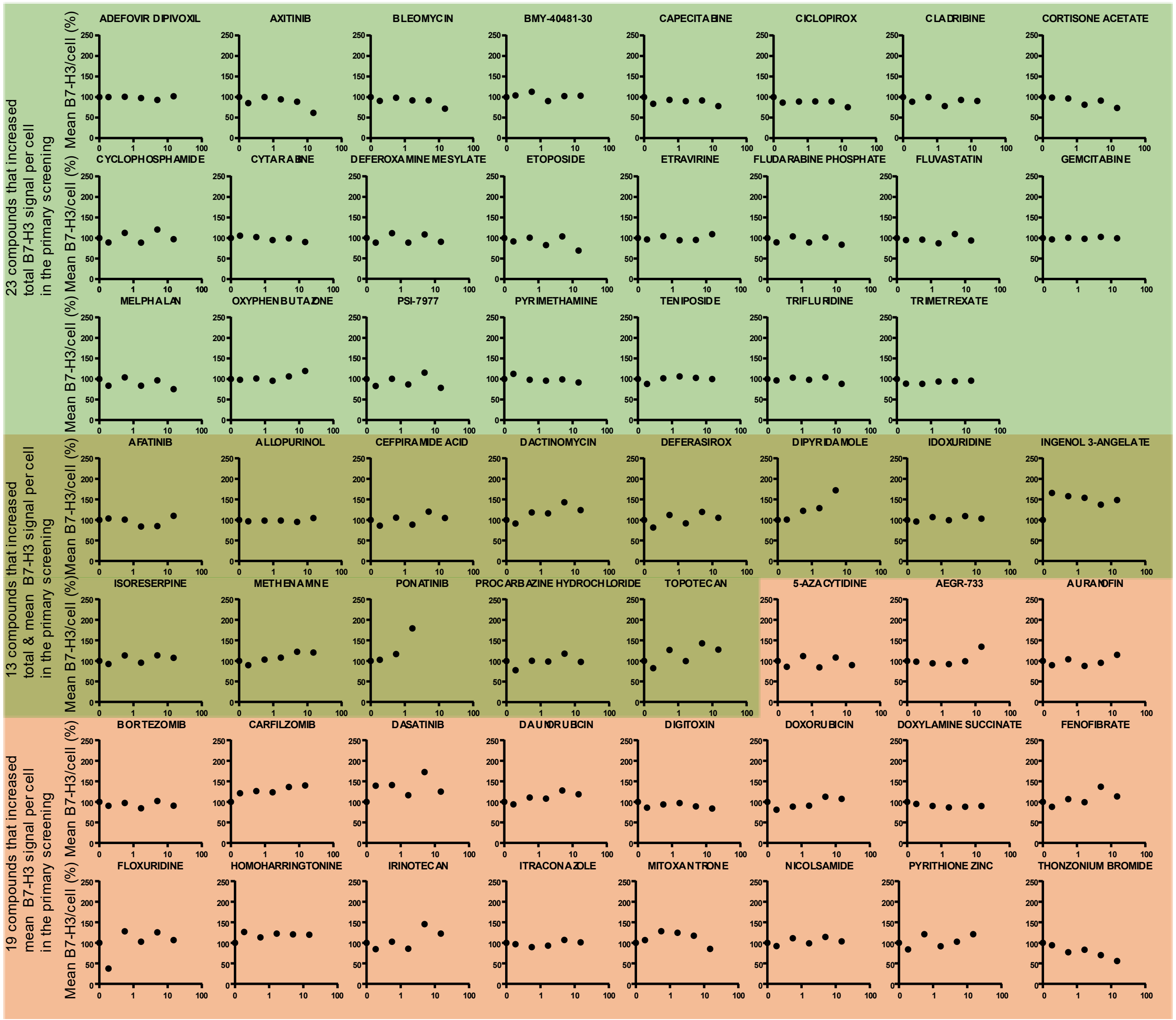

**Figure. S1.**

c. Total B7-H3 signal / cell (%), 48h treatment

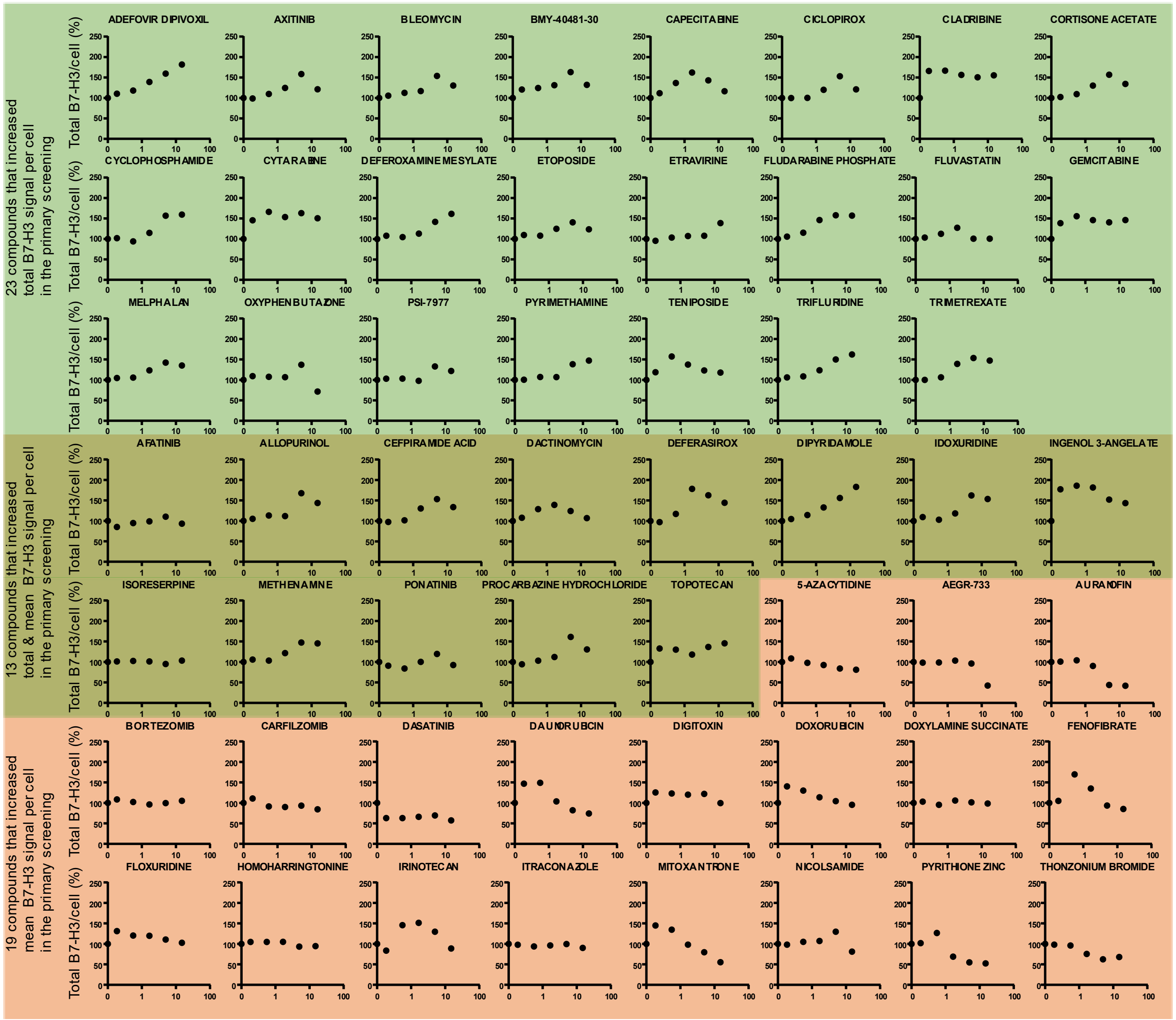

**Figure. S1.**

d. Mean B7-H3 signal / cell (%), 48h treatment

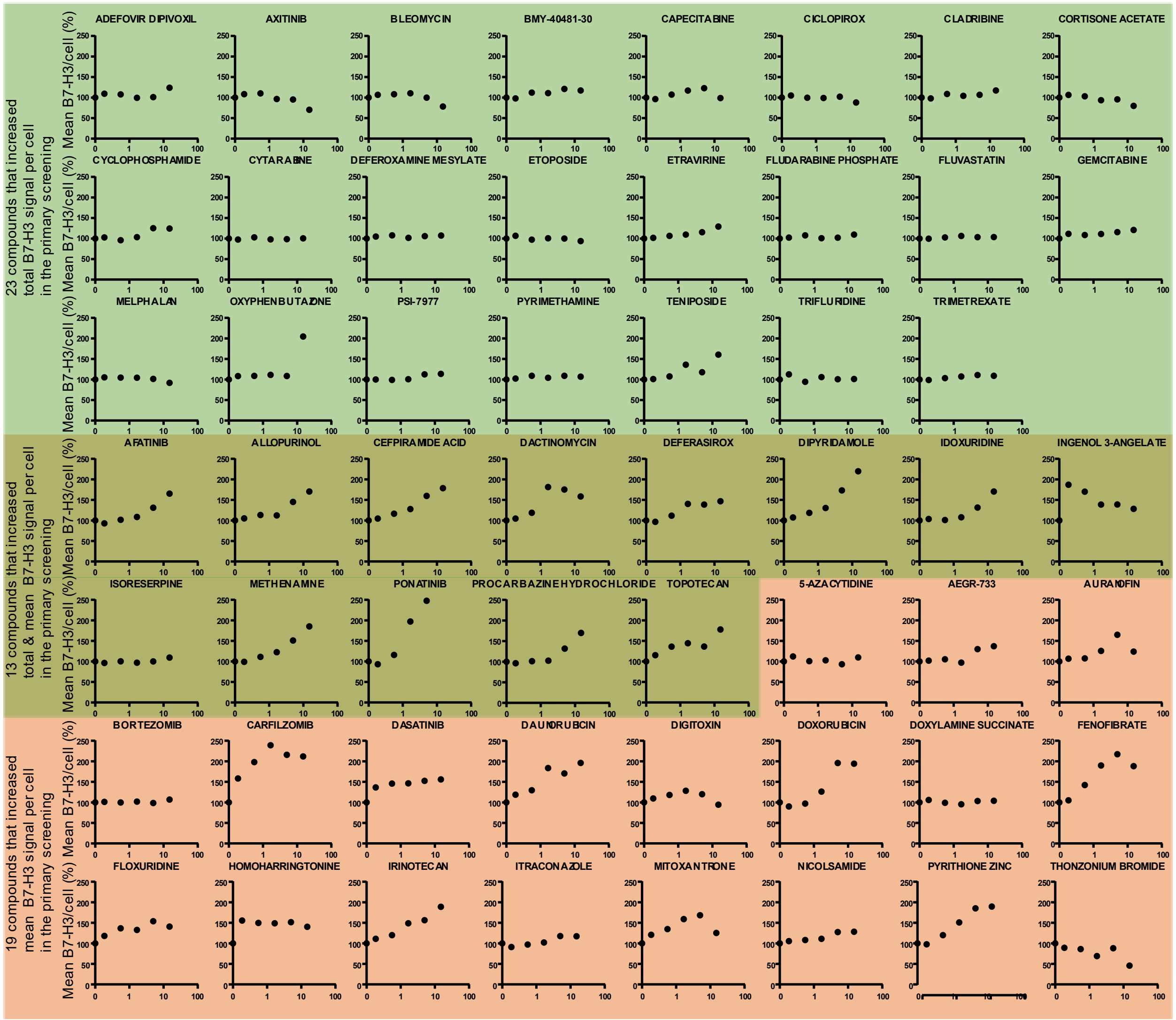


**Figure. S1. Five-point dose-response assay for 55 compounds identified to increase the total and/or mean B7-H3 signal per cell**. Five doses of 55 compounds were treated on LM7 cells for 24 or 48 hours and B7-H3 quantified using the immunofluorescence assay. **a** Total B7-H3 signal per cell after 24 hour treatment with indicated drugs. **b** Mean B7-H3 signal per cell after 24 hour treatment with indicated drugs. **c** Total B7-H3 signal per cell after 48 hour treatment with indicated drugs. **d** Mean B7-H3 signal per cell after 48 hour treatment with indicated drugs. Green background represents compounds that only increased the total B7-H3 signal per cell. Red background represents compounds that only increased the mean B7-H3 signal per cell. Dark green background represents compounds that increased both the total and mean B7-H3 signal per cell.

**Figure. S2.**

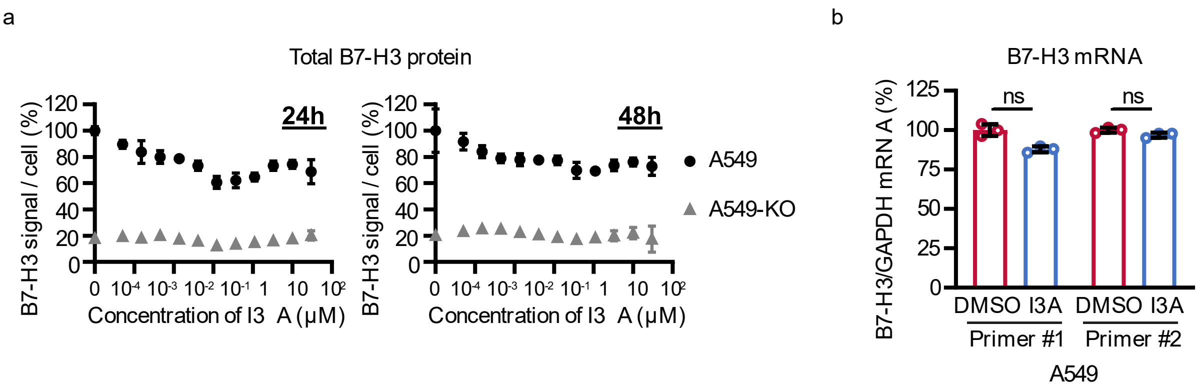


**Figure. S2. Ingenol-3-angelate (I3A) does not increase B7-H3 protein or mRNA expression in A549 cells. a** Total B7-H3 expression per cell in A549 or B7-H3 knockout A549 (A549-KO) cells after treatment with 12 doses of I3A for 24 or 48 hours (N=3). **b** 0.5μM I3A was treated on A549 cells for 48 hours and total RNA was extracted and reverse-transcribed to cDNA. cDNA was analyzed by RT-qPCR. The B7-H3 mRNA level was normalized by the GAPDH mRNA level. Data represents mean ± SD (a and b). ns, non-significant by unpaired t-test (b).

**Figure. S3.**

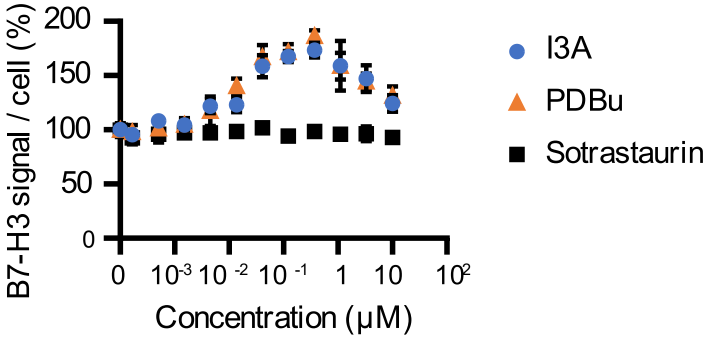


**Figure. S3. The PKC antagonist sotrastaurin does not change B7-H3 protein expression level in LM7 cells.** LM7 cells were treated with 12 doses of I3A, PDBu, or sotrastaurin for 48 hours and B7-H3 protein expression determined by the immunofluorescence assay (N=4). The total B7-H3 signal per cell was normalized by averages of DMSO-treated wells. Data represents mean ± SD.

**Figure. S4.**

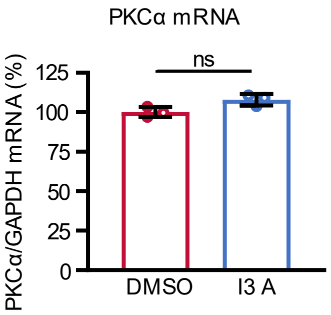


**Figure. S4. I3A does not change the PKCα mRNA level in LM7 cells.** LM7 cells were treated with 0.5 μM I3A or DMSO for 48 hours (N=3). Total RNA was isolated from the cells and reverse-transcribed to cDNA. PKCα mRNA expression level was detected by RT-qPCR. Data represents mean ± SD. ns, non-significant by unpaired t-test.

**Table S1.**

| Compound name | Mechanism | >50% increase | | | |
|---|---|---|---|---|---|
| | | Total B7-H3 signal/cell | | Mean B7-H3 signal/cell | |
| | | 24 hours | 48 hours | 24 hours | 48 hours |
| ADEFOVIR DIPIVOXIL | Reverse-transcriptase inhibitor | | | y | |
| AFATINIB | ErbB family inhibitor | | | | y |
| ALLOPURINOL | Xanthine oxidase inhibitor | | | y | y |
| AURANOFIN | Redox enzyme inhibitor | | | | y |
| AXITINIB | VEGF receptor inhibitor | | | y | |
| BLEOMYCIN | Glycopeptide antibiotic | | | y | |
| BMY-40481-30 | Topoisomerase II inhibitor | | | y | |
| CARFILZOMIB | Proteasome inhibitor | | | | y |
| CEFPIRAMIDE ACID | Penicillin-binding protein | y | | y | y |
| CICLOPIROX | Polyvalent cation chelator | | | y | |
| CLADRIBINE | DNA synthesis and repair inhibitor | | | y | |
| CORTISONE ACETATE | Glucocorticoid receptor binder | | | y | |
| CYCLOPHOSPHAMIDE | Alkylating agent | | | y | |
| DACTINOMYCIN | Transcription inhibitor | | | | y |
| DASATINIB | Src kinase inhibitor | | y | | y |
| DAUNORUBICIN | Topoisomerase II inhibitor | y | | y | y |
| DEFERASIROX | Iron chelator | | | y | |
| DEFEROXAMINE MESYLATE | Iron chelator | | | y | |
| DIPYRIDAMOLE | phosphodiesterase inhibitor | y | y | y | y |
| DOXORUBICIN | Topoisomerase II inhibitor | y | y | y | y |
| FENOFIBRATE | PPAR alpha activator | | | y | y |
| FLOXURIDINE | DNA synthesis inhibition | | | | y |
| FLUDARABINE PHOSPHATE | DNA synthesis inhibition | | | y | |
| GEMCITABINE | DNA synthesis inhibition | | | y | |
| HOMOHARRINGTONINE | Protein synthesis inhibition | | | | y |
| IDOXURIDINE | Antiviral agent | | | y | y |
| INGENOL 3-ANGELATE | Protein kinase C activator | | y | y | y |
| IRINOTECAN | Topoisomerase I inhibitor | | | y | y |
| METHENAMINE | Antimicrobial activity | | | y | y |
| MITOXANTRONE | Topoisomerase II inhibitor | y | | | y |
| OXYPHENBUTAZONE | Cyclooxygenase inhibitor | | | | y |
| PONATINIB | Multi-tyrosine kinase inhibitor | y | y | | y |
| PROCARBAZINE | Alkylating agent | | | y | y |
| PYRITHIONE ZINC | Antifungal effect | | | | y |
| TENIPOSIDE | Topoisomerase II inhibition | | | y | y |
| TOPOTECAN | Topoisomerase I inhibitor | y | | | y |
| TRIFLURIDINE | Viral replication inhibition | | | y | |
| TRIMETREXATE | Dihydrofolate reductase inhibition | | | y | |

**Table. S1. Drugs that increased B7-H3 total or mean B7-H3 expression in the 5-point dose response assay.** Of the 55 drugs evaluated in the 5-point dose-response assay, 38 increased total or mean B7-H3 signal per cell by greater than 50% at one or more doses within 24 or 48 hours. The table includes each of these 38 drug names, mechanisms of action, and the conditions tested. "y" denotes conditions where a drug increased B7-H3 signal per cell by greater than 50%.
